# Integration of metabolism and regulation reveals rapid adaptability to growth on non-native substrates

**DOI:** 10.1101/2022.05.21.492926

**Authors:** Vikas D. Trivedi, Sean F. Sullivan, Debika Choudhury, Venkatesh Endalur Gopinarayanan, Taylor Hart, Nikhil U. Nair

**Author notes:** Corresponding author; <u>@nair_lab</u>; <u>@</u>. Department of Structural Biology and Center for Data Driven Discovery, St. Jude Children’s Research Hospital, Memphis, TN.

## Abstract

Engineering synthetic heterotrophy (i.e., growth on non-native substrates) is key to the efficient bio-based valorization of renewable and waste substrates. Among these, engineering hemicellulosic pentose utilization has been well-explored in *Saccharomyces cerevisiae* (yeast) over several decades – yet the answer to what makes their utilization inherently recalcitrant remains elusive. Through implementation of a semi-synthetic regulon, we find that harmonizing cellular and engineering objectives are key to obtaining highest growth rates and yields with minimal metabolic engineering effort. Concurrently, results indicate that “extrinsic” factors – specifically, upstream genes that direct flux of pentoses into central carbon metabolism – are rate-limiting. We also reveal that yeast metabolism is innately highly adaptable to rapid growth on non-native substrates and that systems metabolic engineering (i.e., flux balancing, directed evolution, functional genomics, and network modeling) is largely unnecessary. We posit that the need for extensive engineering espoused by prior works is a consequence of unfortunate (albeit avoidable) antagonism between engineering and cellular objectives. We also found that deletion of endogenous genes to promote growth demonstrate inconsistent outcomes that are genetic-context- and condition-dependent. For the most part, these knockouts also lead to deleterious pleiotropic effects that decrease the robustness of strains against inhibitors and stressors associated with bioprocessing. Thus, at best, perturbation of “intrinsic” factors (e.g., native metabolic, regulatory genes) results in incremental and inconsistent benefits. At worst, they are detrimental. Overall, this work provides insight into the limitations and pitfalls to realizing efficient synthetic heterotrophy using traditional/systems metabolic engineering approaches, demonstrates the innate adaptability of yeast for metabolism of non-native substrates, and provides an alternate, novel, holistic (and yet minimalistic) approach based on integrating non-native metabolic genes with a native regulon system.

## INTRODUCTION

Engineering metabolism for growth on non-native substrates (i.e., synthetic heterotrophy) has been an outstanding challenge for several decades in various microbial species (*1-4*). The traditional approach is to constitutively overexpress catabolic genes that input substrate to central carbon metabolism with the expectation that the native cellular systems can direct subsequent steps required for growth. In case initial designs do not lead to rapid growth, flux balancing (*5*), adaptive laboratory evolution (ALE) (*6, 7*), functional genomics (*8, 9*), and directed evolution (*6, 10*) approaches are used. More recently, systems-level analysis of regulatory structures in engineered strains have been used to identify dysregulated pathways and corrective interventions have provided avenues to improve engineering outcomes (*11-13*). Unifying to these approaches is that they are reactive interventions that try to rectify inefficiencies introduced due to strain engineering but are not proactive in circumventing undesirable outcomes by harmonizing recombinant activity with native cellular objectives. Typical of this approach is engineering the yeast *Saccharomyces cerevisiae* for complete and rapid utilization of lignocellulosic pentoses (*14-17*). In this work, we engineer strains of yeast for rapid growth and pentose (xylose, arabinose) utilization and assess intrinsic and extrinsic constraints and pitfalls that control desirable phenotypic outcomes. Specifically, we answer the longstanding unresolved question: *what are the inherent/intrinsic limitations within this yeast that prevent it from metabolizing non-native pentose substrates at rates equivalent to native substrates?* Surprisingly, we find that this yeast is intrinsically highly adaptable to rapid growth on non-native substrates and that it can achieve maximum growth rates with minimal engineering – which is in direct contrast with the currently accepted paradigm.

To achieve this, we first modified our semi-synthetic regulon (*18*) – a system based on synthetically activating the galactose (GAL) response system using a non-native substrate – for efficient utilization of arabinose through the isomerase pathway (*19*). We demonstrate that while initial outcomes – like with xylose, previously – are superior to constitutively overexpressing the same genes, growth rates were modest (μ = 0.14 – 0.17 h^-1^). To identify factors that constrain growth on either pentose, we performed pathway balancing, directed evolution, and systems biology-driven functional genomics. To our surprise, we found that when cells can coordinate global growth responses with substrate use through the semi-synthetic GAL regulon (REG), growth is largely extrinsically controlled – i.e., limited by the upstream/heterologous pathway that controls non-native substrate uptake and flux to central carbon metabolism. We also found through parallel investigations that genetic interventions identified through traditional/systems metabolic engineering that prune cellular metabolic and/or regulatory networks to improve strain performance often led to pleiotropic defects with mid-to-severe fitness costs. Thus, our findings suggest that an approach that synergizes activation of the GAL regulon with an optimized upstream heterologous metabolic module reveals the hidden metabolic adaptability of yeast.

## RESULTS

### A semi-synthetic regulon design outperforms a constitutive overexpression strategy

Previously, we demonstrated that coupling activation of galactose (GAL)-responsive regulon to catabolism and growth on non-native substrate, xylose, enabled faster growth and more complete utilization when compared to the constitutive overexpression of the same genes (*18*). This is because GAL regulon activation upregulates growth responsive genes and suppresses starvation responses, in contrast to the traditional constitutive overexpression approaches where the opposite holds true (**Figure 1A**) (*18*). We wanted to assess whether the benefit was unique to xylose or if growth on other substrates, like arabinose, could also benefit from activation of the GAL regulon. Since Gal3p^Syn4.1^ was engineered to activate the GAL regulon in a xylose-dependent manner, we wondered if arabinose could do so as well since the two sugars are structurally similar. We quantified activation of *GAL1p*-*EGFP* by wild-type Gal3p or engineered Gal3p^Syn4.1^ in the presence of native Gal2p and/or engineered Gal2p^2.1^ permease (*18*). We observed that a Gal3p^Syn4.1^ and Gal2p^2.1^ co-expressing strain showed activation on arabinose with low background and high dynamic range (**Figure 1B**). Interestingly, arabinose showed higher activation of *GAL1p*-*EGFP* than xylose, even though both Gal3p^Syn4.1^ and Gal2p^2.1^ were engineered for activity on xylose (**Figure 1C**). Given robust activation, we constructed a semi-integrant strain by integrating accessory genes including, sensor (*GAL3*^*Syn4*.*1*^), transporter (*GAL2*^*2*.*1*^), and transaldolase (*TAL1*) – expressed under GAL promoters – to generate a “REG” (regulon) strain background (**Figure 1D**). Similarly, as control, and to compare the with the traditional engineering approach, we integrated *GAL2*^*2*.*1*^ and *TAL1* under strong constitutive promoters to generate the “CONS” parental strain. We then assessed the growth of both strains on arabinose or xylose after transformation with plasmid-encoded *araBAD* or *XYLA*3-XKS1* catabolic genes, respectively. In ARA-REG and XYL-REG all genes were expressed under GAL-responsive promoters whereas in ARA-CONS and XYL-CONS all genes were under constitutive promoters (**Figure 1D**). On both carbon sources, strains that used the GAL regulon to coordinate substrate assimilation with global metabolism demonstrated higher growth rates (μ: ARA-REG = 0.14 ± 0.04 h^-1^, XYL-REG = 0.17 ± 0.01 h^-1^) compared to those that constitutively overexpressed the same genes (μ: ARA-CONS = 0.06 ± 0.01 h^-1^, XYL-CONS = 0.11 ± 0.01 h^-1^). They also reached higher final cell densities (OD_600_: ARA-REG = 6.01 ± 1.62, ARA-CONS = 1.31 ± 0.04, XYL-REG = 10.8 ± 0.62, XYL-CONS = 5.3 ± 0.49) (**Figure 1E-F**). We then wanted to assess what factors prevented these REG strains from achieving maximum aerobic growth rate (μ = 0.22 – 0.29 h^-1^) (*13, 20*). We explored two potential limitations – i) intrinsic, i.e., native metabolic or regulatory systems downstream of substrate uptake, and ii) extrinsic, i.e., upstream metabolic modules (primarily heterologous genes) that direct non-native substrates to central carbon metabolism.

**Figure 1.**
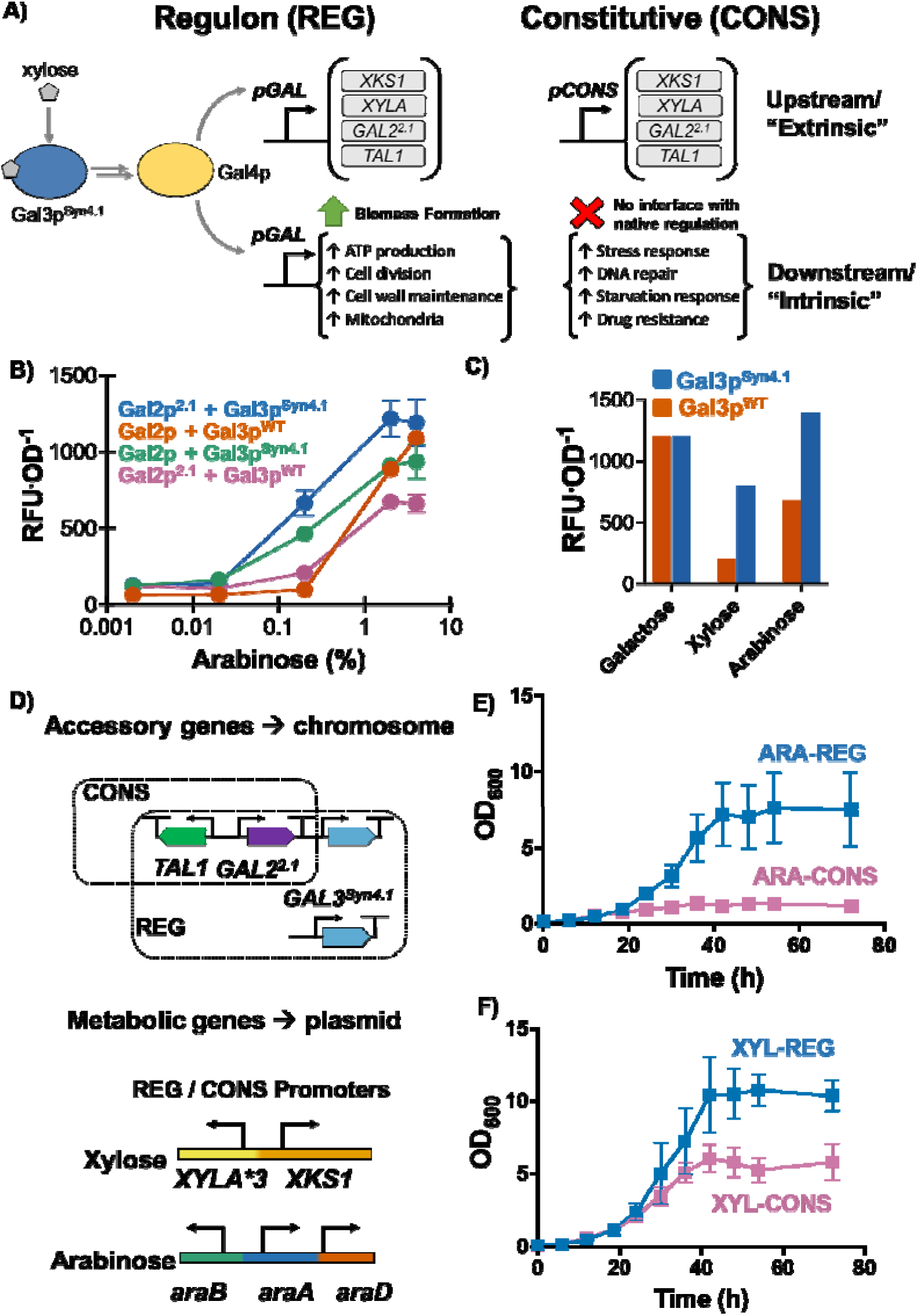
Implementation of a semi-synthetic regulon design for synthetic heterotrophy. **A)** The GAL regulon, synthetically activated by Gal3p^Syn4.1^, regulates GAL-responsive genes as well as hundreds of downstream genes to promote growth. The traditional approach of constitutive overexpression cannot activate downstream genes, which leads to starvation response and slow growth (*18*). **A)** Dose-dependent *GAL1p-EGFP* activation by arabinose of strain expressing Gal3p^WT^ or Gal3p^Syn4.1^ in and Gal2p^WT^ or Gal2p^2.1^. **B)** Comparison of *GAL1p* activation by 2 % of native sugar, galactose, and non-native sugars – xylose and arabinose. **C)** Genetic structure of semi-integrant REG and CONS strain backgrounds. Growth curve of REG and CONS strains on **D)** arabinose and **E)** xylose when transformed with respective arabinose and xylose isomerase pathway genes, respectively. REG and CONS strains use GAL-responsive and constitutive promoters, respectively, for all genes indicated (**Table S1 and S2**).

### Perturbation of intrinsic (downstream) factors result in modest and inconsistent improvements

In traditional/systems metabolic engineering strategies, adaptive laboratory evolution (ALE) or rational data-driven approaches are often used to improve growth rates of initial strain designs, since growth rates and biomass yields are most often suboptimal. We decided to use a (data-driven) systems biology approach to identify “intrinsic” genetic targets to modify to improve the growth rates of our REG strains on xylose and arabinose (i.e., pentoses). By “intrinsic” factors, we mean native yeast genes that are far downstream of substrate uptake and catabolism (e.g., cell wall biosynthesis, DNA repair).

As the GAL regulon has evolved to function harmoniously during growth on a native substrate, we hypothesized that identifying expression differences between galactose and the two non-native substrates and closing the gap between them may aid in better synchronizing GAL regulation with pentose metabolism. To do so, we performed RNA-seq on REG strains grown on arabinose, xylose, and galactose to characterize their respective expression profiles. A differential gene expression (DGE) analysis between individual sugars revealed that a total of 838 genes were differentially regulated between arabinose and galactose and 1,430 genes between xylose and galactose strains (**Figure S1**) (p-value < 0.05 after Benjamini–Hochberg correction). However, rather than pursue the individual substrate comparisons, we chose to repeat the DGE analysis to compare pentose REG vs. galactose utilization to identify “intrinsic” factors that differentiate growth on the two pentoses vs. the hexose galactose. Upon doing so, we found 1,845 genes that were relatively upregulated on pentoses and 1,678 genes that were relatively upregulated during growth on galactose (**Figure 2A**). Given the large number of differentially expressed genes (3,523) between the two conditions (pentoses vs. galactose), we sought to use gene regulatory networks (GRN) to identify highly-connected, regulatory elements that could serve as major contributors to the observed divergence in transcriptional profiles and therefore growth rates. To do so, we first identified three different yeast GRNs that were previously validated for their ability to improve the accuracy of yeast growth phenotypes when integrated with existing metabolic models (*21*). These GRNs – EGRIN, YEASTRACT, and CLR – all sought to capture influences on expression between yeast genes but differed in the data types (gene expression, DNA-binding, protein-protein interactions, etc.) used to infer these connections. The resulting networks were represented as graphs with nodes corresponding to genes and edges between nodes corresponding to an inferred connection between genes. To utilize these networks, we began by associating each node with the expression fold-change and (Benjamini-Hochberg corrected) significance values from the differential expression comparison between the pentose- and galactose-grown REG strains. We then eliminated all nodes with corrected p-values >0.05 along with any connections to those nodes to isolate significantly perturbed sections of the network. We next utilized the network visualization and analysis software tool Cytoscape (*22*) to assign a betweenness centrality (BC), a measure of the centrality of a given node in a network, value to the remaining genes in each network. Plotting BC vs significance in gene expression change helped narrow down a subset of regulatory genes as targets for deletion (**Figure 2B**). Finally, we manually curated a subset of 24 genes among these that have roles in regulation, metabolism, and stress response for deletion to test their effect of growth and phenotype (**Figure 2C**).

**Figure 2.**
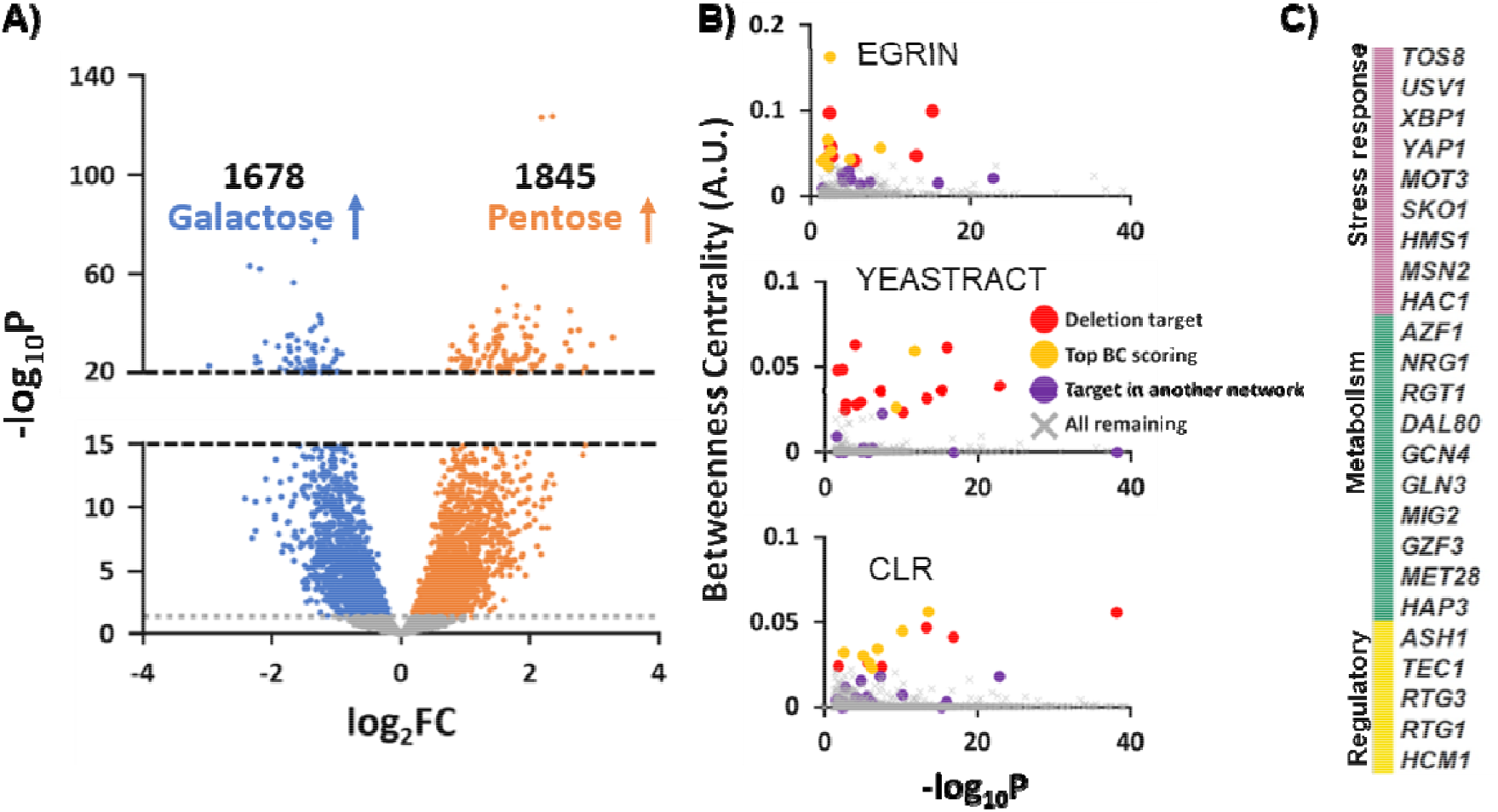
Systems analysis to identify intrinsic limitations in yeast for growth on pentoses. **A)** Differential expression volcano plot for transcriptomic comparison of strains grown individually on galactose or pentose sugars (xylose or arabinose) with the number of genes relatively upregulated in each condition overlaid. **B)** Plots of Betweenness Centrality (BC) (arbitrary units) vs. significance (-log_10_P) for the subset of significantly (p < 0.05) differentially expressed genes demonstrate how native gene targets for deletion were identified using three yeast gene regulatory networks (GRNs): EGRIN (top), YEASTRACT (center), and CLR (bottom). Genes were categorized as either, scoring highest in BC in a given network and picked for deletion (red), not picked for deletion (gold), identified as deletion target in another network (purple), or all remaining significant nodes (grey). **C)** 24 highly connected genes identified via differential gene expression and gene regulatory network analyses (with p < 0.05) as targets for deletion to improve growth on pentoses.

We generated barcoded deletion (KO) libraries of these “intrinsic” factors in wild-type and REG strain backgrounds and performed growth enrichments on glucose, galactose, xylose, and arabinose (**Figure 3A**). Comparing population shifts using barcodes, we identified eight genes that displayed positive fitness on both xylose and arabinose but not on galactose – indicating a role in controlling growth primarily on pentoses. We then tested their growth individually on galactose, xylose, and arabinose (**Figure 3B-D**) and found that only Δ*GLN3* was either neutral or beneficial on both pentoses – all other deletions were detrimental for growth on xylose. Further, the growth rates of all knockout strains were still lower than that of the parental strain on galactose (μ = 0.24 ± 0.03 h^-1^ when *GAL1-7-10* are expressed from a plasmid, **Figure S2**), indicating additional limitations. Overall, the systems metabolic engineering approach failed to identify genetic factors broadly beneficial to growth on pentoses. In contrast, the genome-wide expression changes effected by GAL regulon activation are beneficial for all three substrates.

**Figure 3.**
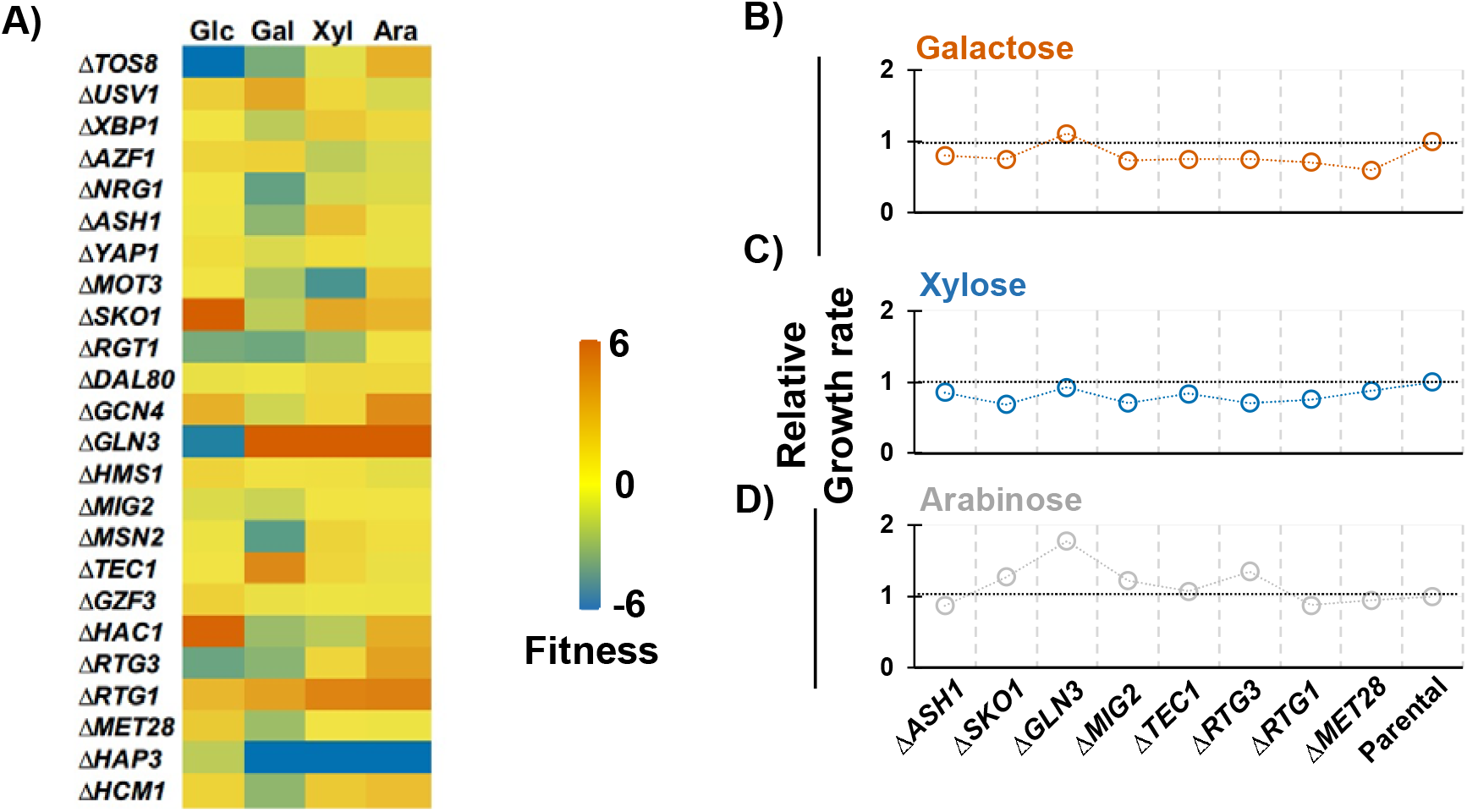
Effect of gene knockouts on growth phenotype. **A)** Fitness map of knockout library of 24 genes identified from the mutual information network. The scale represents fitness ranging from blue (negative) to orange (positive). Growth rates of KOs with positive fitness on **B)** galactose, **C)** xylose (*GAL1p-XYLA*3 GAL10p -XKS1*), **D)** arabinose (*GAL1p-araB GAL10p-araA GAL7p-araD*). Glc = glucose, Gal = galactose, Xyl = xylose, Ara = arabinose. The growth rates were normalized from the average of at least two biological replicates.

### Pentose metabolism is largely extrinsically (upstream) controlled with a regulon approach

Our studies have so far focused on identifying “intrinsic” factors that may be controlling/bottlenecking growth on pentoses and given the inconsistent benefits, we wondered whether the limitations were, in fact, “extrinsic”. The former implies that the native metabolic and/or regulatory capacity of this yeast is inherently limited for effective pentose metabolism and improving growth rate requires vast restructuring of associated (intrinsic) metabolic and/or regulatory networks. The latter implies that there are no inherent limitations in this yeast and that it is already poised for rapid growth on pentoses, but the observed low growth rates are due to suboptimal design of the upstream metabolic module that includes the heterologous (extrinsic) genes. To assess whether the “extrinsic limitation” paradigm has any merits, we needed to optimize the design of the upstream metabolic module responsible for substrate uptake and flux into central carbon metabolism (i.e., glycolysis).

First, we looked at the effect of plasmid copy number. For arabinose, there was no difference in growth rate when *araA-araB-araD* genes were expressed on high- or low-copy plasmids, whereas for xylose, we only observed growth when the *XYLA*3-XKS1* gene dose was high (**Figure S3**). In either case, changing plasmids backbone did not improve growth rate. Next, we hypothesized that balancing expression of these heterologous genes may be required to enhance growth rate. For arabinose, we created all six combinations of gene-promoter pairings and assessed their performance. While such promoter swaps have been previously demonstrated to improve strain performance (*23*), we were surprised at the marked improvement in growth rate – from 0.14 ± 0.04 h^-1^ in the original design to 0.27 ± 0.03 h^-1^ in the best re-design (**Figure 4A**). To understand the cause of this behavioral change, we compared the relative expression levels of the *araA, araB*, and *araD* using quantitative reverse transcription PCR (qPCR) (**Figure 4B**). We observed that in the poor performing combinations, expression of *araB* (ribulokinase) was the highest, whereas, in high performing combinations, expression level followed this pattern *araA* > *araB* > *araD*. In addition, we found a strong positive correlation between growth rate and relative expression of *araA*:*araB* and *araA*:*araD*, respectively (**Figure 4C-E**). The success of this approach encouraged us to attempt the same on xylose and we found similar improvements – from 0.17 ± 0.01 h^-1^ to 0.24 ± 0.01 h^-1^ (**Figure S4**). These are comparable to the growth rate of yeast on galactose when *GAL1-7-10* are expressed from a plasmid (0.24 ± 0.03 h^-1^) (**Figure S2**).

**Figure 4.**
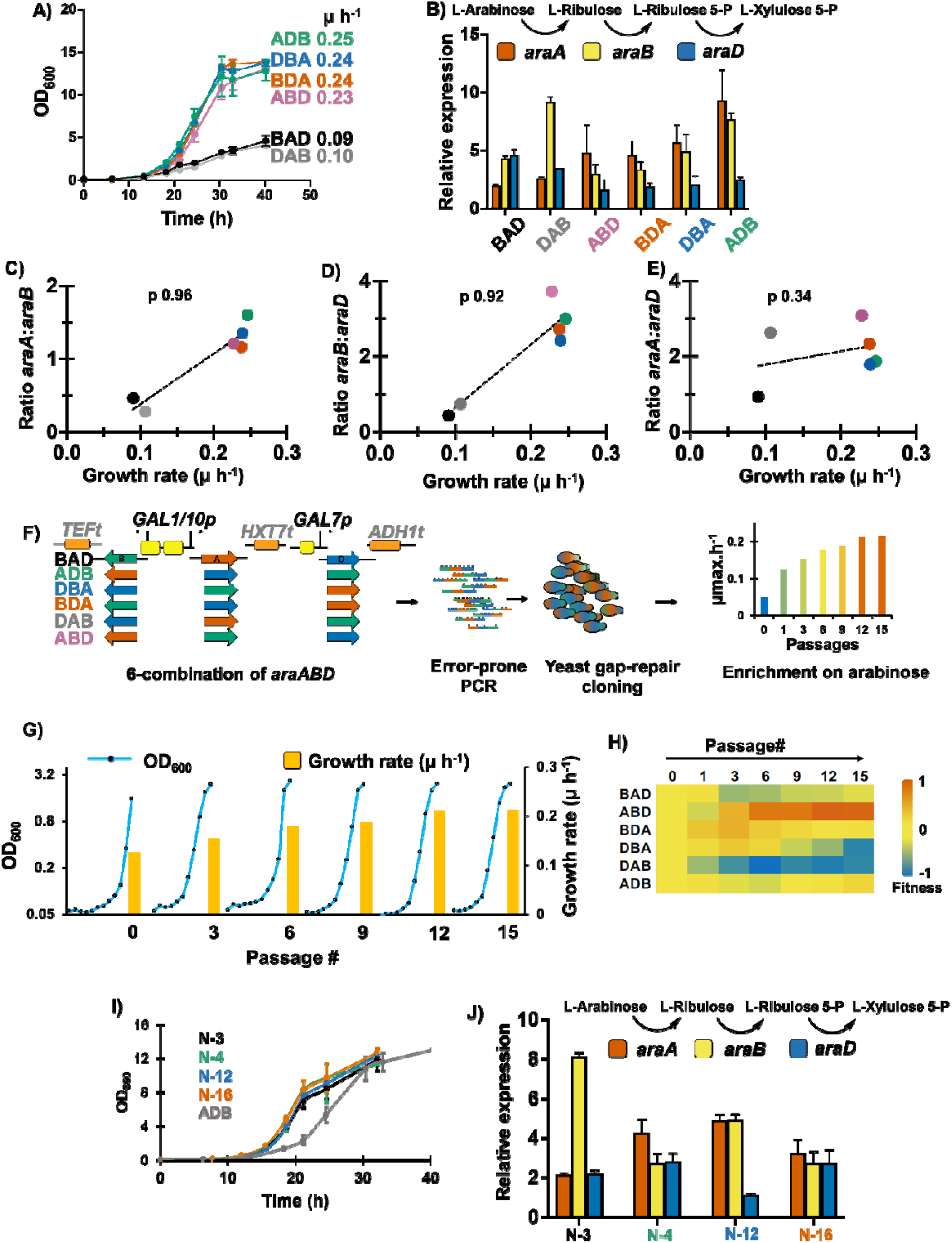
Optimizing extrinsic (upstream) factors to enhance growth of yeast on pentoses. **A)** Comparison of growth performance of different promoter-gene combination. **B)** Gene expression analysis of the six strains by qPCR. Pearson correlation coefficient between growth rate and gene expression levels of **C)** *araA*:*araB*, **D)** *araB:araD*, and **E)** *araA:araD*, respectively. **G)** Enrichment of mutagenic *araBAD* library in SC+Ara. **H)** Fitness heatmap of strain dynamics during enrichment on arabinose, revealed by sequencing barcodes. **I)** Growth curves of top variants from the directed evolution study with parental control strain (ADB) as reference, and **J)** qPCR expression profile of *araBAD* genes encoded in plasmids isolated from top performing strains isolated after directed evolution.

Finally, we used directed evolution to improve the growth rate on arabinose further (**Figure 4F**). We randomly mutagenized the six arabinose pathway combinations, adding barcodes to track the lineages and enriched this library size of 10^8^ variants in minimal (SC+Ara) and complex (2YP+Ara) arabinose media (**Figure 4F**). Over the course of 15 subcultures, the growth rate over subcultures increased from 0.12 h^-1^ to 0.22 h^-1^ and 0.18 h^-1^ to 0.26 h^-1^ in SC+Ara and 2YP+Ara, respectively (**Figure 4G, S6**). Using barcodes, we tracked the performance of promoter-gene combinations (**Figure 4H**). We observed that initially all the six plasmids start at similar abundance, but the *araA-B-D* (under *GAL1-10-7p*, respectively) was the most abundant at the end of enrichment in SC+Ara (**Figure 4H**). We picked single colonies from each condition and calculated their growth rates and identified 4 variants that showed the highest growth rates (0.35 ± 0.04 h^-1^) from the SC+Ara enriched culture (**Figure S5**). We sequenced the barcodes to identify the lineage and the whole cassette to identify the mutations and found that three of the six initial designs were represented in the four best variants (named N-3, N-6, N-12, and N-16) (**Table S3**). Re-transformation into parent background strain indicated that the four variants attained the same maximum growth rate as that on galactose (0.29 ± 0.01 h^-1^), which is the highest reported growth rate on any pentose reported in the literature for this yeast (**Figure 4I**). Since the mutations were distributed throughout the cassettes, we quantified the expression levels of *araA, araB*, and *araD* in the four strains (**Figure 4J**) and found patterns that differed significantly than parental strains (**Figure 4B**). We saw a strong positive correlation between growth rate and relative expression of *araA*:*araB* and negative correlation between *araB*:*araD*, indicating that the directed evolution campaign likely altered both activity and expression in each strain differently (**Figure S6**). These results highlight a key insight about yeast: it’s ability to utilize pentoses is largely limited only by “extrinsic” factors (upstream pathway, especially heterologous enzyme activity) and minimally by any “intrinsic” factor (i.e., native regulation or metabolic pathway).

### Engineering intrinsic (downstream) genes lead to pleiotropic fitness trade-offs

Next, we assessed whether the global regulatory elements identified through our network analysis would benefit our best designs (*araA-B-D* under *GAL1-10-7p* and *XYLA*3-XKS1* under *GAL1-7p*) but found that deletion of each of the 8 genes did not result in any improvements over the parental strains (**Figure 5A**). We considered exploring combinatorial deletions and ALE to attempt to further enhance growth rates on these two substrates. However, we expected that significant improvements were unlikely as both strains had already attained rates comparable to the empirically determined maximum aerobic growth rates of this yeast (0.22 h^-1^ – 0.29 h^-1^) (*13, 20*) with high biomass yields and short lag, we expect that improvements would be insignificant.

**Figure 5.**
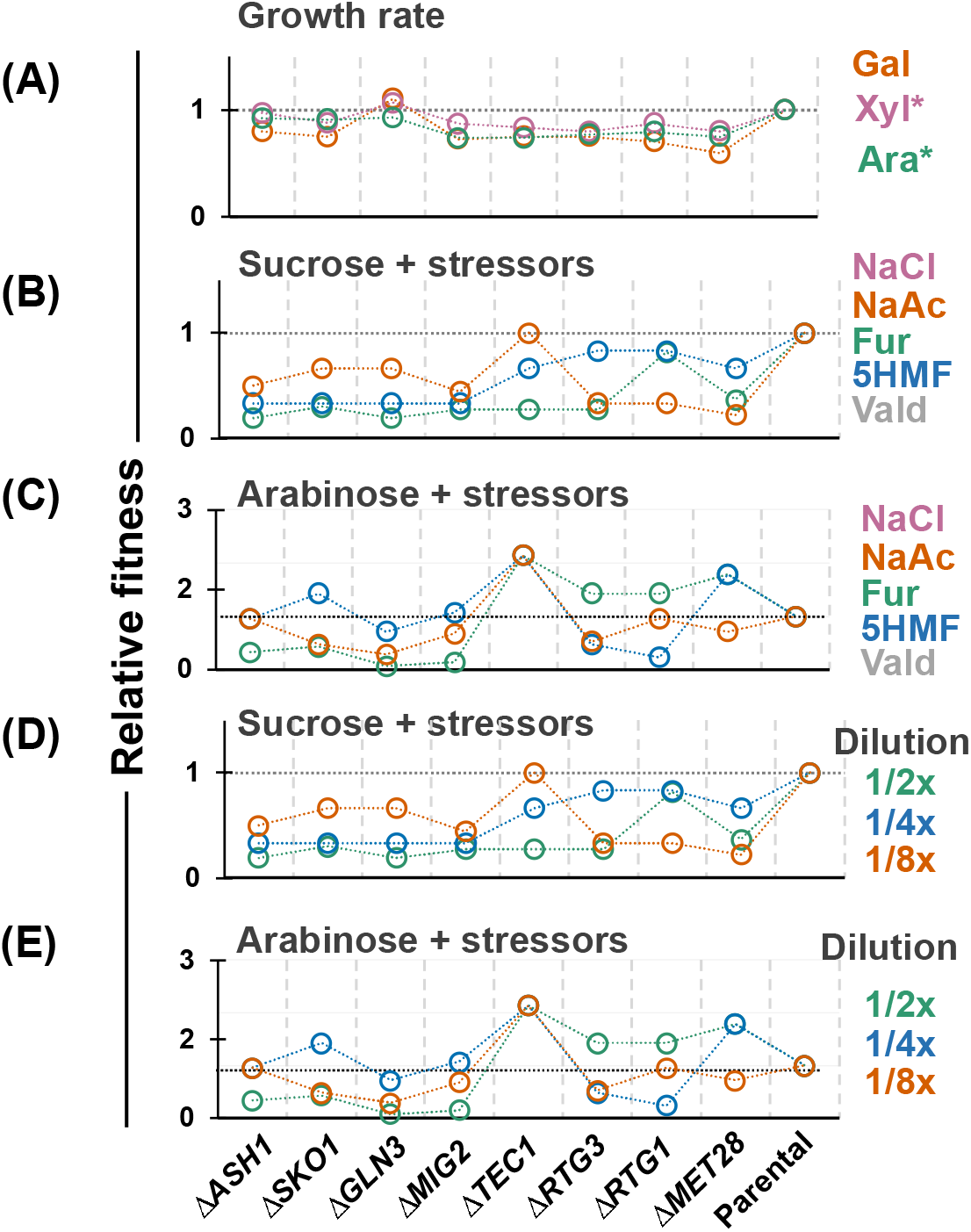
Effect of gene deletions on growth and stress-response phenotypes. **A)** Growth rate of the optimized plasmids in KO-strains. Fitness of KOs in presence of individual stressors when grown on **B)** sucrose and **C)** arabinose or in the mixture of stressors on **D)** sucrose and **E)** arabinose. The relative fitness is expressed as growth of deletion strains over the fitness of parental strain under the same conditions. NaCl = sodium chloride; NaAc = sodium acetate; Fur = furfural; 5HMF = 5-hydroxymethylfurfural; Vald = veratraldehyde. 1/2x, 1/4x, 1/8x = serial dilutions of the five stressors combined. Xyl* = *GAL10p-XYLA*3 GAL7p-XKS1*; Ara* = *GAL1p-araA GAL10p-araB GAL7p-araD* in respective REG backgrounds.

One consideration not yet assessed is the suitability of these strains for bioprocessing and their resilience to growth inhibitors associated with bioprocessing applications. Given that all deletion targets are highly-connected genes that control key cellular processes, we were concerned that dysregulation major networks may lead to undesirable pleiotropic effects. Since resilience against stress is a complex phenotype, often requiring concerted response from gene networks, we tested the fitness of all single knockout (KO) strains on several stressors (sodium chloride, sodium acetate, furfural, 5-hydroxymethylfurfural, and veratraldehyde) relevant to bioprocessing individually and in combination (**Figure 5B-E**). Using parental strain as reference, we observed that performance of strains under stress was highly context-dependent. For example, in sucrose medium, all deletions lost fitness in single or mixed inhibitor cultures (**Figure 5C, E**). Conversely, on arabinose (with GAL regulon activated), certain KO strains had improved tolerance to stressors (e.g., Δ*TEC1*, Δ*MET28*) (**Figure 5D, F**). Δ*GLN3* has previously been shown to improve fitness under isobutanol stress (*24*); however, in our study, it was less fit than the parental strain under all stress conditions. Interestingly, it did demonstrate improved growth rate in a sub-optimal upstream metabolic design in arabinose (**Figure 3E**) but lost that benefit in a more optimized design (**Figure 5C, E**). Collectively, these results highlight that deleting genes to remodel expression profiles to enhance a single phenotype (e.g., growth rate on a non-native substrate) can lead to some improvements, but they are often accompanied by negative pleiotropic effects that make the strain less suitable for eventual bioprocessing applications where growth conditions are often non-ideal.

## DISCUSSION

Despite decades of effort engineering synthetic heterotrophy with *S. cerevisiae*, there is no formalized understanding of what limits the metabolic adaptability of this yeast for growth on pentoses. It is important to differentiate between substrate utilization/uptake from growth since the former can be readily achieved by diverting flux to unwanted or dead-end byproducts (e.g., Crabtree metabolites, organic acids, pentitols) often with native substrate supplementation to support biomass generation. However, valorization to high-value products requires efficient catabolism to central metabolic products whose concentrations and fluxes are regulated and associated with growth. Initial studies in literature focused on upstream (“extrinsic”) elements (e.g., heterologous gene expression/activity, transporter engineering, etc. (*25-29*)) whereas recent focus has been more “intrinsic”, focusing on functional genomics, adaptively laboratory evolution (ALE), and often network/systems analysis, to identify downstream limitations (*5-13*). Successes have been abundantly reported with improvements in growth along with a series of deletion and overexpression targets (*PHO13, ALD6, ASK10, YPR1, SNF6, RGT1, CAT8, MSN4, GPD1, CCH1, ADH6, BUD21, ALP1, ISC1, RPL20B, COX4, ISU1, SSK2, YLR042c, CYC8, PHD1, TEC1, ARR1*, etc. (*13, 30-36*)). Significantly, many targets are unique to individual studies. This is perhaps unsurprising as the impact of such interventions, and even the performance of identical heterologous pathways, can depend on genetic context (*37, 38*). Typifying the traditional approach is a recent report where all the aforementioned extensive systems metabolic engineering approaches were used to develop a strain of yeast with high growth aerobic growth rate (0.26 h^-1^) and short lag phase (9-15 h) on xylose (*13*). Despite this, it is not clear if the insights were translatable to engineering growth on any other substrate.

Through our work, we posit that the requirement for extensive engineering to identify intrinsic limitations is only required when the initial strain design is sub-optimal. Indeed, constitutive overexpression of upstream metabolic genes results in significant stress (*18*) and low growth rates that must be compensated for through systems approaches focused on preventing cellular detection of stress rather than activating growth promoting systems. We demonstrated that with an optimal upstream module, strains can attain superior growth profiles (fast specific growth rate, μ ≥ 0.24 h^-1^; short lag phase, <10 h; high final biomass, OD_600_ >10) if a growth-associated regulon – the GAL regulon – is activated. This builds on our prior observation that synthetic activation of the GAL regulon potentiates cells for rapid growth on non-native substrates by upregulating growth responsive genes and suppressing starvation responses (*18*). Importantly, using this approach our final strain designs had minimal modifications – we deleted only one gene (other than the Leloir pathway genes) – *GRE3* – to minimize oxidation of substrate pentoses to pentitols (*39*). Many genes that have previously been identified as important inactivation targets to improve growth on pentoses are intact. We also overexpressed only a minimal upstream metabolic module (*TAL1* and *GAL2*^*2*.*1*^) along with the specific pentose metabolic genes (*araBAD* or *XYLA*3-XKS1*). Our work strongly argues that growth on pentoses is largely extrinsically controlled and there are no major intrinsic regulatory or metabolic limitations in this yeast. This insight presents a paradigm shift in engineering synthetic heterotrophy.

An additional advantage of this approach is the preservation of native regulatory systems that are otherwise dysregulated in the traditional engineering approach. We found that while deletion of endogenous genes could improve growth in strains with sub-optimal upstream modules under certain conditions, they are associated with pleiotropic defects and are less robust in the presence of growth inhibitors. This is not surprising since most genetic interventions aim to suppress cellular responses to stress rather than promote growth. Thus, strains engineered for synthetic heterotrophy through extensive gene inactivation are less robust for eventual bioprocessing applications. This highlights a major drawback of traditional/systems metabolic engineering approaches like ALE. In contrast, our regulon-based approach is minimalistic and holistic – our strains maintain native regulatory systems and even exploit them toward the engineering goal (i.e., the GAL regulon). On the one hand, our utilization of a single strain background (W303-1a) and catabolic pathway for each substrate in this study could potentially limit the broad applicability of our findings. For example, the use of the oxidoreductase xylose pathway has been demonstrated to improve resistance to lignocellulosic inhibitors relative to the isomerase pathway (*40*). Also, quantitative measures of strain fitness (i.e., growth rates) are known to be influenced by seemingly confounding factors like the choice of selection marker and the copy number of the plasmid bearing the auxotrophy-complementing gene (*41*). However, the regulon approach we are advocating is not an isolated perturbation to the cell but interfaces with a native regulatory system that coordinates the expression of hundreds of genes. Thus, we believe it is more likely to be transferable to different strain backgrounds and target substrates. Furthermore, this approach significantly simplifies and expedites the design-build-test cycle for synthetic heterotrophy and demonstrates the intrinsic adaptability of yeast toward growth on non-native substrates. We expect that this insight and approach will expand the utility of this yeast for valorizing current and emerging waste and/abundant substrates.

## DATA AVAILABILITY

RNA-seq sequencing data has been submitted to NCBI SRA and is available under accession PRJNA837644. R script for mutual information network is available at https://github.com/nair-lab/yeast-MINet.

## ACKNOWLEDGEMENTS

The authors would like to thank current and former Nair lab members, Dr. Todd C. Chappell, and Dr. Karishma Mohan for helpful discussions. This work was supported by NIH grant #DP2HD91798, NSF grant #1935354, and Tufts Launchpad | Accelerator to N.U.N.

## AUTHOR CONTRIBUTIONS

N.U.N., V.D.T., and D.C. conceived and designed the research project. V.D.T., S.F.S., D.C., and N.U.N. co-wrote the manuscript. V.D.T., S.F.S., V.E.G., and T.H. performed the experiments. V.D.T., S.F.S., and N.U.N. analyzed the data. All the authors have reviewed the manuscript and approved it for submission.

## MATERIALS AND METHODS

### Strains and plasmids

The list of plasmids and strains used are listed in Supplementary **Table S1 and S2**. Strain W303-1a (*MATa leu2-3,112 trp1-1 can1-100 ura3-1 ade2-1 his3-11,15*) and plasmids pIS374, pIS376, pIS385 were obtained from Euroscarf (Germany). The plasmids were constructed in the present using NEB-HiFi DNA assembly master mix from NEB (Beverly, MA).

### Growth studies

Overnight inoculums were grown in the required dropout SC medium (6.7 g/L yeast nitrogen base without amino acids, 2 g/L dropout mix) with sucrose (2 %). The culture was washed twice in the growth medium and resuspended at an initial OD_600_ of 0.1 with appropriate sugar (2 %) in 250□mL shake flasks containing 20□mL of media. OD_600_ measurement was checked at frequent time intervals (3-6 h) on SpectraMax M3 spectrophotometer (Molecular Devices). Growth rate was determined by plotting the values in GraphPad Prism following non-linear regression and using exponential growth equation, *Y = Y*_*0*_ *exp(kX)*.

### Directed evolution

Error prone PCR libraries of arabinose cassettes (pVDT14, pVDT38-42) were generated as described earlier (*18*). Six-barcoded primers (**Table S4**) were used to track the proportion of the lineage during the enrichment. The amplicons from error prone PCR were assembled into pRS413 plasmids using yeast gap-repair cloning. To attain high library size, the linear fragments were transformed into yeast by electroporation as described previously (*42*). We attained library size of ∼10^8^ CFUs which was then subjected to enrichment on arabinose in 2YP (20 g/L yeast extract, 40 g/L peptone, 100 mg/L adenine hemisulfate) as well SC media. The cell pellets from each passage were frozen for quantifying barcodes using amplicon sequencing (Genewiz, Cambridge, MA). At the end of enrichment, the library pool was plated on 2YP+Ara and SC+Ara and 18 colonies were randomly picked for growth rate determination. The plasmids from best variants were isolated from strain, transformed into *E. coli*. The corresponding plasmid isolated from *E. coli* was sequenced and re-transformed into REG-strain for growth rate determination.

### Genomics integrations

Accessory cassettes (pVDT29, 30, 35, **Table S1**) were integrated into VEG16 (W303-1a, Δ*GRE3*, Δ*GAL1*, Δ*GAL3*, Δ*GAL7*, Δ*GAL10*) (*18*) strain via disintegrator plasmid (*43*). Counterselection was performed on SC medium with 1□g/L 5-FOA to remove the *URA3* marker and the locus was amplified, and sequence confirmed.

### Deletion library

To generate the deletion library, the cassettes for the required targets were amplified from the genomic DNA of the appropriate strains from the KO collection (*44*). Since the cassettes contained *KANMX* marker, the library was selected on YP+G418 (400 μg/mL). The pool was stored as a stock and used for studying fitness by transforming it with plasmids for galactose, xylose, and arabinose utilizing cassettes.

### qPCR expression analysis

Total yeast RNA isolation was performed on OD ∼ 1 of yeast cells collected from mid-log phase growth using the GeneJet RNA Purification Kit (Thermofisher Scientific Catalog #: K0731) according to the instructions for isolating yeast RNA. Two samples were collected for each growth condition. The concentration of total RNA isolated was obtained by measuring 10-fold dilutions from each sample on a spectrophotometer. Using these concentrations, 2.5 μg of total RNA was subjected to the ‘routine’ DNase treatment protocol using the Invitrogen Turbo DNA-free Kit (Thermofisher Scientific Catalog #: AM1907). Next, 20 % of the final reaction (10-50 μL, targeting 500 ng RNA) was used to synthesize cDNA for the sample using the Invitrogen SuperScript IV First-Strand Synthesis System (Thermofisher Scientific Catalog #: 18091050). The random hexamers supplied by the kit were used to prime the reverse transcriptase enzyme. cDNA samples were diluted 100-fold (2 μL into 198 μL dH_2_O), of which 2 μL was used as the template for a 15 μL qPCR reaction using the Applied Biosystems PowerUp SYBR Green Master Mix (Thermofisher Scientific Catalog #: A25741). Each sample was tested for the expression level of five genes: the three arabinose catabolic genes (*araA, araB*, and *araD*) as well as two housekeeping genes (*TFC1* and *UBC6*) using the primers listed in **Table S5**. Reactions were run on an Applied Biosystems Quantstudio 5 Real-Time PCR Instrument. The relative expression level of each catabolic genes was calculated by subtracting the geometric mean of the cycle thresholds (Ct) of the two housekeeping genes from the Ct of that catabolic gene, followed by conversion of the log-base two Ct into a non-log value that can be compared across genes.

### RNA-seq

Transcriptomics of strains WT, XYL-REG, ARA-REG were performed on the mid-log phase cultures grown on their respective carbon source (galactose, xylose, or arabinose). Cells pellets were washed twice in water and stored at −80□°C and outsourced to Genewiz Inc. for RNA extraction and sequencing. RNA-seq was performed on Illumina HiSeq. Raw FASTQ files were processed for differential expression analysis using GeneiousPrime. The possible adapter sequences and low-quality short-read (less than 50_bp) trimming were performed using BB-Trim package. The reads were aligned to the reference genome W303 obtained from Saccharomyces Genome Database (http://www.yeastgenome.org) using Bow-Tie2 package. The edgeR package was used to normalize the gene count based on library size and was converted to cpm (counts per million) using. DESEQ2 package was sued for differential gene expression analysis. Genes with p-values < 0.05 and fold change of ≥ 2 were considered as differentially expressed.

### Amplicon sequencing analysis

The total DNA was extracted and quantified using a spectrophotometer and approximately 100 ng was used as template for PCRs. Unique, barcoded primers flanked by Illumina sequencing adaptors were used to generate the amplicons after 15-20 cycles of PCR for sequencing (**Table S6**). The barcoded samples were pooled and sent for amplicon sequencing (2×250 bp, Genewiz, New Jersey, USA). For each pooled sample, we received approximately 100,000 sequencing reads. These data were processed according to a previously described bioinformatic workflow using Geneious Prime® 2020.2.4 (*45*). Briefly, the reads were paired and merged using the BBMerge package and filtered for poor-quality reads using the BBDuk package. The reads were mapped using BowTie2 and gene expression was calculated and differential expression was determined using DESeq2 (*46*).

### Network analysis

The *S. cerevisiae* genes that we consider as the ‘gold standard’ list of transcription factors is the unique set of genes identified in the file supplemental data **File S1**. Our analysis utilizes three yeast genetic regulatory networks (EGRIN, YEASTRACT, and CLR) (*21*) in the context of integrating regulatory and metabolic networks to enable more accurate prediction of yeast phenotypes in different growth conditions (*47*). The Environment and Gene Regulatory Influence Network (EGRIN) consists of 92 regulators and 2,588 interactions and was constructed using the cMonkey and Inferelator computational tools trained on expression data from 2,929 microarray experiments to identify gene clusters and potential regulatory genes. The YEASTRACT gene regulatory network was extracted directly from the YEASTRACT database2 and consists of 177 regulators and 31,075 regulatory associations based on both ‘direct evidence’ – chromatin immunoprecipitation (ChIP), ChIP-on-chip, electrophoretic mobility shift assay, or examination of the effect of TF binding site mutations on target gene expression – as well as ‘indirect evidence’ provided by gene expression changes in response to deletion, mutation, or overexpression of a given TF3.

The EGRIN, YEASTRACT, and CLR yeast gene regulatory networks (GRN) were downloaded as Microsoft Excel files from the supplementary information of a recent study by Wang et al. (*21*). These excel files captured the GRN as two-column lists of genes with each row representing an inferred connection between the genes. We imported these files as networks in the open-source network visualization and analysis software Cytoscape (*22*). For each network, we created spreadsheets that associates the fold-change in expression and corrected p-value from the differential gene expression analysis between the pentose (REG) and galactose grown strains with each unique yeast gene found in that network. By importing these files into Cytoscape, we were able to map this additional information onto the nodes in the networks. Next, each network was filtered by removing all nodes corresponding to genes with Benjamini-Hochberg corrected p-values > 0.05. The “Analyze Network” function was used to generate betweenness centrality (BC), a quantitative measure of the centrality of a given node in a network, values for each of the remaining nodes. The final list of 24 deletion targets was manually curated from a combined list of the top 15 genes by BC from each of the three networks based on which were most likely to be directly related to the metabolism of pentoses. The R programming language was used for importing and manipulating data. R scripts can be found in https://github.com/nair-lab/yeast-MINet.

### Stressor studies

We determined fitness of the strains in presence of various stressors encountered during biomanufacturing using plate assay. Briefly, we tested the growth of by performing spot dilution on 2YP supplemented with 2 % sucrose or arabinose and various concentration of the stressors (**Table S7**). The plates after incubation were imaged and the colonies were counted to determine the CFUs. The CFUs in the presence of the stressor was divided by the CFUs in absence of the stressor to determine the fitness of that strain. This value was normalized to the fitness of the parental strain and was expressed as the relative fitness.

## REFERENCES

1. J. Sun, W. Sun, G. Zhang, B. Lv, C. Li, High efficient production of plant flavonoids by microbial cell factories: Challenges and opportunities. Metabolic Engineering, (2022).

2. M. A. Franden et al., Engineering Pseudomonas putida KT2440 for efficient ethylene glycol utilization. Metabolic engineering 48, 197–207 (2018).

3. S. Pontrelli et al., Escherichia coli as a host for metabolic engineering. Metabolic engineering 50, 16–46 (2018).

4. T. Sun, Y. Yu, K. Wang, R. Ledesma-Amaro, X.-J. Ji, Engineering Yarrowia lipolytica to produce fuels and chemicals from xylose: A review. Bioresource Technology 337, 125484 (2021).

5. T. J. Hanly, M. A. Henson, Dynamic flux balance modeling of microbial co-cultures for efficient batch fermentation of glucose and xylose mixtures. Biotechnology and bioengineering 108, 376–385 (2011).

6. S.-M. Lee, T. Jellison, H. S. Alper, Directed evolution of xylose isomerase for improved xylose catabolism and fermentation in the yeast Saccharomyces cerevisiae. Applied and Environmental Microbiology 78, 5708–5716 (2012).

7. L. Diao et al., Construction of fast xylose-fermenting yeast based on industrial ethanol-producing diploid Saccharomyces cerevisiaeby rational design and adaptive evolution. BMC biotechnology 13, 1–9 (2013).

8. D. J. Wohlbach et al., Comparative genomics of xylose-fermenting fungi for enhanced biofuel production. Proceedings of the National Academy of Sciences 108, 13212–13217 (2011).

9. A. Matsushika, H. Inoue, T. Kodaki, S. Sawayama, Ethanol production from xylose in engineered Saccharomyces cerevisiae strains: current state and perspectives. Applied microbiology and biotechnology 84, 37–53 (2009).

10. M. Lee, H. J. Rozeboom, E. Keuning, P. de Waal, D. B. Janssen, Structure-based directed evolution improves S. cerevisiae growth on xylose by influencing in vivo enzyme performance. Biotechnology for Biofuels 13, 1–18 (2020).

11. Z. Dai, J. Nielsen, Advancing metabolic engineering through systems biology of industrial microorganisms. Current opinion in biotechnology 36, 8–15 (2015).

12. K. V. Presnell, H. S. Alper, Systems metabolic engineering meets machine learning: A new era for data-driven metabolic engineering. Biotechnology journal 14, 1800416 (2019).

13. X. Li et al., Metabolic network remodelling enhances yeast’s fitness on xylose using aerobic glycolysis. Nature Catalysis 4, 783–796 (2021).

14. L. Sun, Y. S. Jin, Xylose assimilation for the efficient production of biofuels and chemicals by engineered Saccharomyces cerevisiae. Biotechnology Journal 16, 2000142 (2021).

15. X. Wang, J. Yang, S. Yang, Y. Jiang, Unraveling the genetic basis of fast l-arabinose consumption on top of recombinant xylose-fermenting Saccharomyces cerevisiae. Biotechnology and Bioengineering 116, 283–293 (2019).

16. R. Garcia Sanchez et al., Improved xylose and arabinose utilization by an industrial recombinant Saccharomyces cerevisiae strain using evolutionary engineering. Biotechnology for biofuels 3, 1–11 (2010).

17. H. W. Wisselink, M. J. Toirkens, Q. Wu, J. T. Pronk, A. J. van Maris, Novel evolutionary engineering approach for accelerated utilization of glucose, xylose, and arabinose mixtures by engineered Saccharomyces cerevisiae strains. Applied and environmental microbiology 75, 907–914 (2009).

18. V. Endalur Gopinarayanan, N. U. Nair, A semi-synthetic regulon enables rapid growth of yeast on xylose. Nature communications 9, 1–12 (2018).

19. H. W. Wisselink et al., Engineering of Saccharomyces cerevisiae for efficient anaerobic alcoholic fermentation of L-arabinose. Applied and Environmental Microbiology 73, 4881–4891 (2007).

20. Z. Dai, M. Huang, Y. Chen, V. Siewers, J. Nielsen, Global rewiring of cellular metabolism renders Saccharomyces cerevisiae Crabtree negative. Nature communications 9, 1–8 (2018).

21. Z. Wang et al., Combining inferred regulatory and reconstructed metabolic networks enhances phenotype prediction in yeast. PLoS computational biology 13, e1005489 (2017).

22. P. Shannon, Markiel a, Ozier O, et al. cytoscape: a software environment for integrated models of biomolecular interaction networks. Genome Res 13, 1 (2003).

23. C. E. Costa, P. Carvalho, L. Domingues, Strategic combination of different promoters in lactose metabolisation and host chassis selection for high bioethanol titres from dairy wastes. Bioresource Technology Reports 19, 101131 (2022).

24. K. Kuroda et al., Critical roles of the pentose phosphate pathway and GLN3 in isobutanol-specific tolerance in yeast. Cell Systems 9, 534-547. e535 (2019).

25. E. Young, S.-M. Lee, H. Alper, Optimizing pentose utilization in yeast: the need for novel tools and approaches. Biotechnology for biofuels 3, 1–12 (2010).

26. E. M. Young, A. D. Comer, H. Huang, H. S. Alper, A molecular transporter engineering approach to improving xylose catabolism in Saccharomyces cerevisiae. Metabolic engineering 14, 401–411 (2012).

27. J. G. Nijland, A. J. Driessen, Engineering of pentose transport in Saccharomyces cerevisiae for biotechnological applications. Frontiers in Bioengineering and Biotechnology, 464 (2020).

28. B. Wiedemann, E. Boles, Codon-optimized bacterial genes improve L-arabinose fermentation in recombinant Saccharomyces cerevisiae. Applied and environmental microbiology 74, 2043–2050 (2008).

29. V. Endalur Gopinarayanan, N. U. Nair, Pentose metabolism in Saccharomyces cerevisiae: the need to engineer global regulatory systems. Biotechnology journal 14, 1800364 (2019).

30. S. R. Kim et al., Rational and evolutionary engineering approaches uncover a small set of genetic changes efficient for rapid xylose fermentation in Saccharomyces cerevisiae. PloS one 8, e57048 (2013).

31. J. Usher et al., Chemical and Synthetic Genetic Array Analysis Identifies Genes that Suppress Xylose Utilization. (2011).

32. Y. Shen et al., An efficient xylose-fermenting recombinant Saccharomyces cerevisiae strain obtained through adaptive evolution and its global transcription profile. Applied Microbiology and Biotechnology 96, 1079–1091 (2012).

33. L. V. Dos Santos et al., Unraveling the genetic basis of xylose consumption in engineered Saccharomyces cerevisiae strains. Scientific Reports 6, 1–14 (2016).

34. N. S. Parachin, O. Bengtsson, B. Hahn-Hägerdal, M. F. Gorwa-Grauslund, The deletion of YLR042c improves ethanolic xylose fermentation by recombinant Saccharomyces cerevisiae. Yeast 27, 741–751 (2010).

35. J. Zha, M. Shen, M. Hu, H. Song, Y. Yuan, Enhanced expression of genes involved in initial xylose metabolism and the oxidative pentose phosphate pathway in the improved xyloseutilizing Saccharomyces cerevisiae through evolutionary engineering. Journal of Industrial Microbiology and Biotechnology 41, 27–39 (2014).

36. C. F. Wahlbom, R. R. Cordero Otero, W. H. van Zyl, B. r. Hahn-Ha□gerdal, L. J. Jo□nsson, Molecular analysis of a Saccharomyces cerevisiae mutant with improved ability to utilize xylose shows enhanced expression of proteins involved in transport, initial xylose metabolism, and the pentose phosphate pathway. Applied and environmental microbiology 69, 740–746 (2003).

37. J. T. Cunha, T. Q. Aguiar, A. Romaní, C. Oliveira L. Domingues, Contribution of PRS3, RPB4 and ZWF1 to the resistance of industrial Saccharomyces cerevisiae CCUG53310 and PE-2 strains to lignocellulosic hydrolysate-derived inhibitors. Bioresource Technology 191, 7–16 (2015).

38. A. Romaní, F. Pereira, B. Johansson, L. Domingues, Metabolic engineering of Saccharomyces cerevisiae ethanol strains PE-2 and CAT-1 for efficient lignocellulosic fermentation. Bioresource technology 179, 150–158 (2015).

39. K. Traff, R. Otero Cordero, W. Van Zyl, B. Hahn-Hagerdal, Deletion of the GRE3 aldose reductase gene and its influence on xylose metabolism in recombinant strains of Saccharomyces cerevisiae expressing the xylA and XKS1 genes. Applied and environmental microbiology 67, 5668–5674 (2001).

40. J. T. Cunha, P. O. Soares, A. Romaní, J. M. Thevelein, L. Domingues, Xylose fermentation efficiency of industrial Saccharomyces cerevisiae yeast with separate or combined xylose reductase/xylitol dehydrogenase and xylose isomerase pathways. Biotechnology for Biofuels 12, 1–14 (2019).

41. J. T. Pronk, Auxotrophic yeast strains in fundamental and applied research. Applied and environmental microbiology 68, 2095–2100 (2002).

42. J. R. Thompson, E. Register, J. Curotto, M. Kurtz, R. Kelly, An improved protocol for the preparation of yeast cells for transformation by electroporation. Yeast 14, 565–571 (1998).

43. I. Sadowski, T. C. Su, J. Parent, Disintegrator vectors for single-copy yeast chromosomal integration. Yeast 24, 447–455 (2007).

44. G. Giaever et al., Functional profiling of the Saccharomyces cerevisiae genome. nature 418, 387–391 (2002).

45. V. D. Trivedi et al., In-Depth Sequence–Function Characterization Reveals Multiple Pathways to Enhance Enzymatic Activity. ACS Catalysis 12, 2381–2396 (2022).

46. M. I. Love, W. Huber, S. Anders, Moderated estimation of fold change and dispersion for RNA-seq data with DESeq2. Genome biology 15, 1–21 (2014).

47. M. J. Herrgård, B.-S. Lee, V. Portnoy, B. Ø. Palsson, Integrated analysis of regulatory and metabolic networks reveals novel regulatory mechanisms in Saccharomyces cerevisiae. Genome research 16, 627–635 (2006).

