## Supplementary Information for "Integration of metabolism and regulation reveals rapid adaptability to growth on non-native substrates"

**Table S1.** List of plasmids used.

| **Plasmids** | **Description** |
| --- | --- |
| pVEG8-WT | *pRS426, 2μ URA3 ADH1t-EGFP-GAL1p/GAL10p-KANMX-HXT7t-GAL3p-GAL3^WT^-TEF1t* |
| pVEG8-Syn4.1 | *pRS426, 2μ URA3 ADH1t-EGFP-GAL1p/GAL10p-KANMX-HXT7t-GAL3p-GAL3^Syn4.1^-TEF1t* |
| pVEG11 (REG) | *pRS426, 2µ URA3 ADH1t-Piromyces_XYLA*3-GAL1p/GAL10p-XKS1-HXT7t* |
| pVEG15 (CONS) | *pRS426, 2µ URA3 ADH1t-Piromyces_XYLA*3-TEF1p-TPI1p-XKS1-HXT7t* |
| pVDT14 | *pRS416, CEN URA3 ADH1t-araB-GAL1p/GAL10p-araA-HXT7t-GAL7p-araD-TEF1t* |
| pVDT23 (REG) | *pRS426, 2µ URA3 ADH1t-araB-GAL1p/GAL10p-araA-HXT7t-GAL7p-araD-TEF1t* |
| pVDT22 (CONS) | *pRS426, 2µ URA3 ADH1t-araB-TEF1p-TPI1p-araA-HXT7t-GPM1p-araD-TEF1t* |
| pVDT29 | *pIS385, URA3 GAL3p-GAL3^Syn4.1^-TEF1t-ADH1t-GAL2^2.1^-GAL1/10p-TAL1-HXT7t* |
| pVDT30 | *pIS376, URA3 GAL1p-GAL3^Syn4.1^-GAL3t* |
| pVDT35 | *pIS385, URA3 ADH1t-GAL2^2.1^-TEF1p/TPI1p-TAL1-Hxt7t* |
| pVDT38 | *pRS416, CEN URA3 ADH1t-araA-GAL1p/GAL10p-araB-HXT7t-GAL7p-araD-TEF1t* |
| pVDT39 | *pRS416, CEN URA3 ADH1t-araB-GAL1p/GAL10p-araD-HXT7t-GAL7p-araA-TEF1t* |
| pVDT40 | *pRS416, CEN URA3 ADH1t-araD-GAL1p/GAL10p-araB-HXT7t-GAL7p-araA-TEF1t* |
| pVDT41 | *pRS416, CEN URA3 ADH1t-araD-GAL1p/GAL10p-araA-HXT7t-GAL7p-araB-TEF1t* |
| pVDT42 | *pRS416, CEN URA3 ADH1t-araA GAL1p/GAL10p-araD-HXT7t-GAL7p-araB-TEF1t* |
| pVDT48 | *pRS413, CEN HIS3 ADH1t-Piromyces_XYLA*3-GAL1p/GAL10p -XKS1-HXT7t* |
| pVDT49 | *pRS413, CEN HIS3 ADH1t-Piromyces_XYLA*3-TEF1p-TPI1p-XKS1-HXT7t* |
| pVDT53 | *pRS426, 2µ URA3 GAL1t-GAL1-GAL1p/GAL10p-GAL10-GAL10t-GAL7p-GAL7-GAL7t* |
| pVDT54 | *pRS423, 2µ HIS3 GAL1t-GAL1-GAL1p/GAL10p-GAL10-GAL10t-GAL7p-GAL7-GAL7t* |
| pVDT55 | *pRS423, 2µ HIS3 ADH1t-Piromyces_XYLA*3-GAL1p/GAL10p-XKS1-HXT7t* |
| pVDT56 | *pRS423, 2µ HIS3 ADH1t- araB- GAL1p/GAL10p -araA-HXT7t-GAL7p-araD-TEF1t* |
| pVDT57 | *pRS423, 2µ HIS3 ADH1t-XKS1-GAL10p/GAL1p -TEF1t -GAL7p-Piromyces_XYLA*3-HXT7t* |
| pVDT58 | *pRS423, 2µ HIS3 ADH1t-XKS1-GAL1p/GAL10p-TEF1t-GAL7p-Piromyces_XYLA*3-HXT7t* |
| pVDT59 | *pRS423, 2µ HIS3 ADH1t-XKS1-GAL1p/GAL10p-Piromyces_XYLA*3-HXT7t* |
| pVDT60 | *pRS423, 2µ HIS3 ADH1t-XKS1-GAL10p/GAL1p-Piromyces_XYLA*3-HXT7t* |
| pVDT61 | *pRS423, 2µ HIS3 ADH1t-XKS1-GAL7p-TEF1t-GAL1p/GAL10p -Piromyces_XYLA*3-HXT7t* |
| pVDT62 | *pRS423, 2µ HIS3 ADH1t-XKS1-GAL7p-TEF1t-GAL10p/GAL1p-Piromyces_XYLA*3-HXT7t* |

**Table S2**. List of strains used.

| **Strain** | **Description** |
| --- | --- |
| W303-1a | *MATa LEU2-3,112 TRP1-1 CAN1-100 URA3-1 ADE2-1 HIS3-11,15* |
| VEG16 | *W303-1a ΔGAL3 ΔGRE3 ΔGAL1 ΔGAL7 ΔGAL10* |
| VEG20 | *VEG16 GAL2p-GAL2^2.1^-TEF1t::leu2* |
| VDT13 | *VEG16 GAL2^2.1^-GAL1p-GAL10p-TAL1-GAL3p-GAL3^Syn4.1^::lys2*  *GAL1p-GAL3^Syn4.1^::leu2* |
| VDT27 | *VEG16 ADH1t-GAL2^2.1^-TEF1p/TPI1p-TAL1-HXT7t::lys2* |
| VDT53 | *VDT13 ΔASH1::KANMX* |
| VDT54 | *VDT13 ΔSKO1::KANMX* |
| VDT55 | *VDT13 ΔGLN3::KANMX* |
| VDT56 | *VDT13 ΔTEC1::KANMX* |
| VDT57 | *VDT13 ΔRTG3::KANMX* |
| VDT58 | *VDT13 ΔRTG1::KANMX* |
| VDT59 | *VDT13 ΔMET28::KANMX* |
| VDT60 | *VDT13 ΔMIG2::KANMX* |

**Table S3.** Variants identified from directed evolution of arabinose metabolic plasmid.

| **Plasmids** | **Description** |
| --- | --- |
| N-3 (pVDT65) | *pRS413 CEN HIS3 ADH1t-araD*(E8G-K6E)-GAL1p/GAL10p-araA*(K72M-N410Y-A422D-G467A)-HXT7t-GAL7p-araB*(N277Y)-TEF1t* |
| N-4 (pVDT66) | *pRS413 CEN HIS3 ADH1t- araA*(A40S-D398G)-GAL1p/GAL10p-araB*(I441T)-HXT7t-GAL7p-araD*(N232K)-TEF1t* |
| N-12 (pVDT67) | *pRS413 CEN HIS3 ADH1t-araA*(H225R)-GAL1p/GAL10p-araD-HXT7t-GAL7p-araB*(Y56H-A124S-E333V)-TEF1t* |
| N-16 (pVDT69) | *pRS413 CEN HIS3 ADH1t-araA*(E94V)-GAL1p/GAL10p-araB*(F133L-E490G)-HXT7t-GAL7p-araD*(T116S-E208G)-TEF1t* |

**Table S4.** Barcoded primers used to generate error-prone PCR library of cassettes for arabinose utilization.

| **Primer name** | **Sequence (5’→3’)*** |
| --- | --- |
| 14-BC-FP | ACTCACTATAGGGCGAATTG*TACTGCAG*GTCGACTGGATGGCGGCGTTAG |
| 38-BC-FP | ACTCACTATAGGGCGAATTG*GGACTCCT*GTCGACTGGATGGCGGCGTTAG |
| 39-BC-FP | ACTCACTATAGGGCGAATTG*TAGGCATG*GTCGACTGGATGGCGGCGTTAG |
| 40-BC-FP | ACTCACTATAGGGCGAATTG*TAAGGCGA*GTCGACTGGATGGCGGCGTTAG |
| 41-BC-FP | ACTCACTATAGGGCGAATTG*CGTACTAG*GTCGACTGGATGGCGGCGTTAG |
| 42-BC-FP | ACTCACTATAGGGCGAATTG*TATCCTCT*GTCGACTGGATGGCGGCGTTAG |
| ARA-CMN-RP | CAATTAACCCTCACTTGCCGGTAGAGGTGTGGTC |

*Red italics are barcodes.

**Table S5.** Primers used for gene expression studies.

| **Primer name** | **Sequence (5’→3’)** |
| --- | --- |
| TFC1-F | GCGGTATTGACAGCAGGTTCAAA |
| TFC1-R | CCATCCAGTTAGTTCATTCGCCTTA |
| UBC6-F | ACCATCAGAAGAAGACATTAGCAAGA |
| UBC6-R | TCATCACCTGTATTTGCCGCAT |
| *araD*F_RT | AGCCTCATTAGTCAGCCAGCAC |
| *araD*R_RT | AGGCACTTGCACCATGCTTAC |
| *araA*F_RT | CTCACAACAAGCAAACCGC |
| *araA*R_RT | CGAGCAGGATCATCTTTACCC |
| *araB*F_RT | GGACGTTATGATTGCACAGG |
| *araB*R_RT | TGCAGTTTGCCGACAAGC |

**Table S6**. Primers for amplicon sequencing of arabinose enrichment library.

| **Primer name** | **Sequence (5’→3’)** |
| --- | --- |
| Ara-AmpEz-F-1 | ACACTCTTTCCCTACACGACGCTCTTCCGATCT*AGCGATTACG*ATTAAGTTGGGTAACGC |
| Ara-AmpEz-F-2 | ACACTCTTTCCCTACACGACGCTCTTCCGATCT*AGGAATTGCG*ATTAAGTTGGGTAACGC |
| Ara-AmpEz-F-3 | ACACTCTTTCCCTACACGACGCTCTTCCGATCT*GGCTCAACCG*ATTAAGTTGGGTAACGC |
| Ara-AmpEz-F-4 | ACACTCTTTCCCTACACGACGCTCTTCCGATCT*GTTGATGGCG*ATTAAGTTGGGTAACGC |
| Ara-AmpEz-F-5 | ACACTCTTTCCCTACACGACGCTCTTCCGATCT*AGTTATACCGA*TTAAGTTGGGTAACGC |
| Ara-AmpEz-F-6 | ACACTCTTTCCCTACACGACGCTCTTCCGATCT*ACATTCCACGA*TTAAGTTGGGTAACGC |
| Ara-AmpEz-F-7 | ACACTCTTTCCCTACACGACGCTCTTCCGATCT*GGCTATGTCGA*TTAAGTTGGGTAACGC |

*Red italics indicate the barcode.

**Table S7**: Concentration of stressors used in the study.

| **Stressor** | **Concentration** |
| --- | --- |
| 5-Hydroxymethyl furfural (5HMF) | 20 mM |
| Veratraldehyde (Val) | 10 mM |
| Sodium acetate (NaAc) | 100 mM |
| Furfural (Fur) | 10 mM |
| Sodium chloride (NaCl) | 0.5 M |
| Mixture* | 5HMF (1 mM), Val (0.25 mM), NaAc (100 mM), Fur (2.5 mM),  NaCl (0.25 M) |

* Serial dilutions made from this “1x” stock.

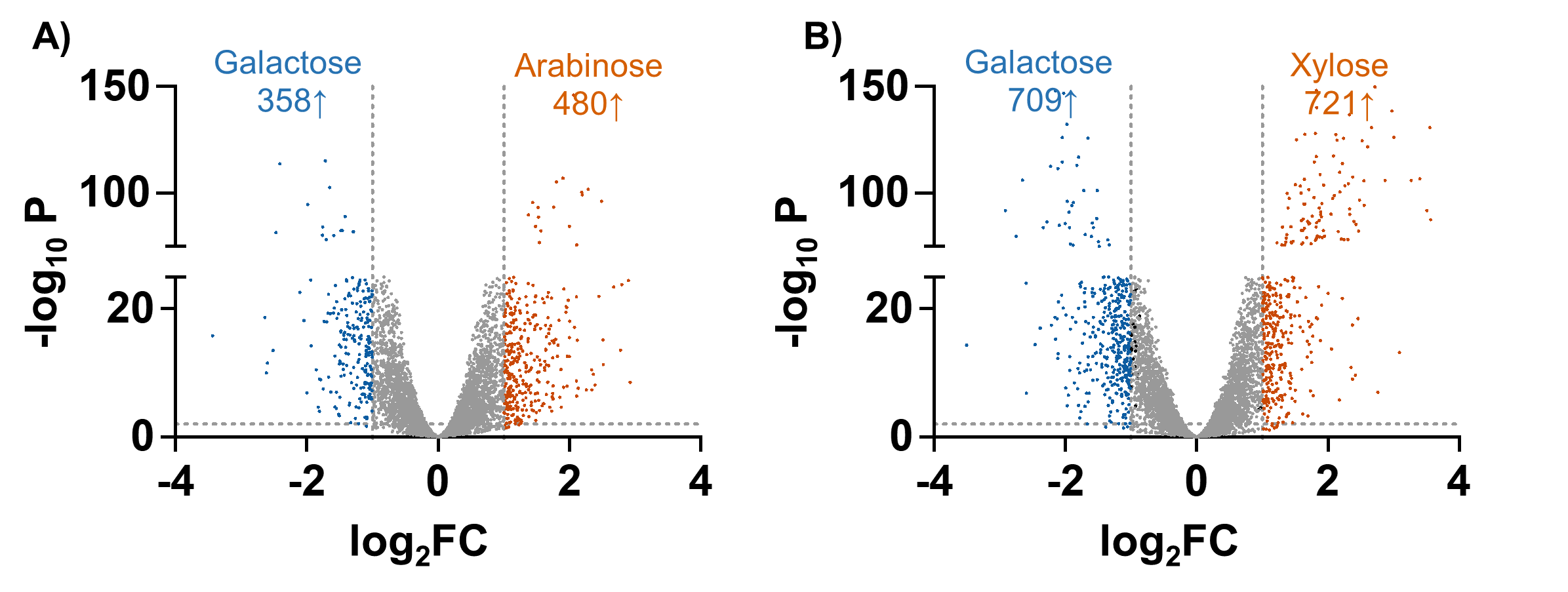

**Figure S1.** Volcano plot of differentially expressed genes in **A)** ARA-REG when compared to galactose. Orange represents the genes upregulated (480) whereas blue represents genes downregulated (358) in arabinose. **B)** XYL-REG when compared to galactose. Orange represents the genes upregulated (721) whereas blue represents genes downregulated (709) in xylose.

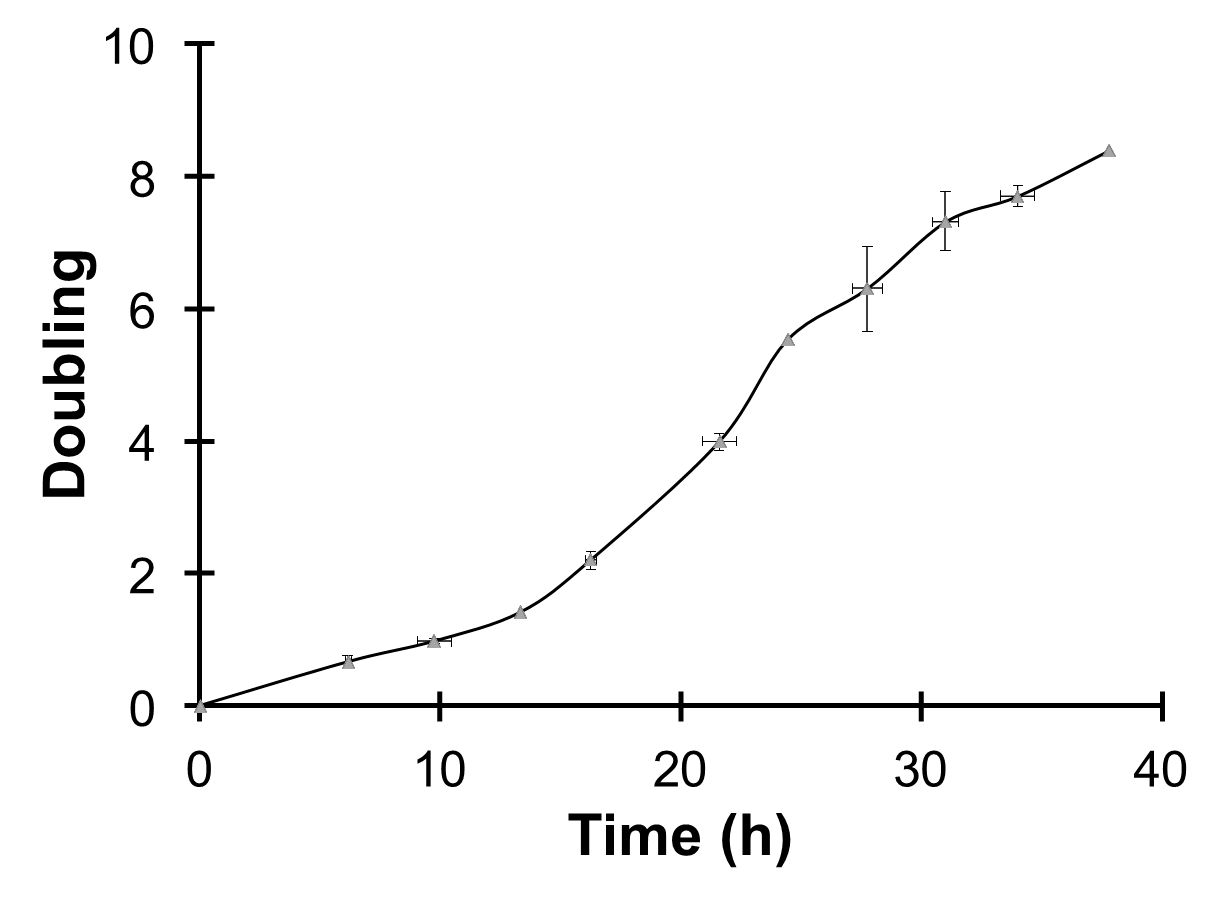

µ = 0.24 h^-1^

**Figure S2.** Growth curve of a REG strain carrying galactose cassette for *GAL1, GAL7, GAL10* on plasmid (pVDT54) under their native promoters. Chromosomal copies of *GAL1-7-10* are deleted. The culture grown on SC media with sucrose (2%) was inoculated in 2YP with galactose (2%).

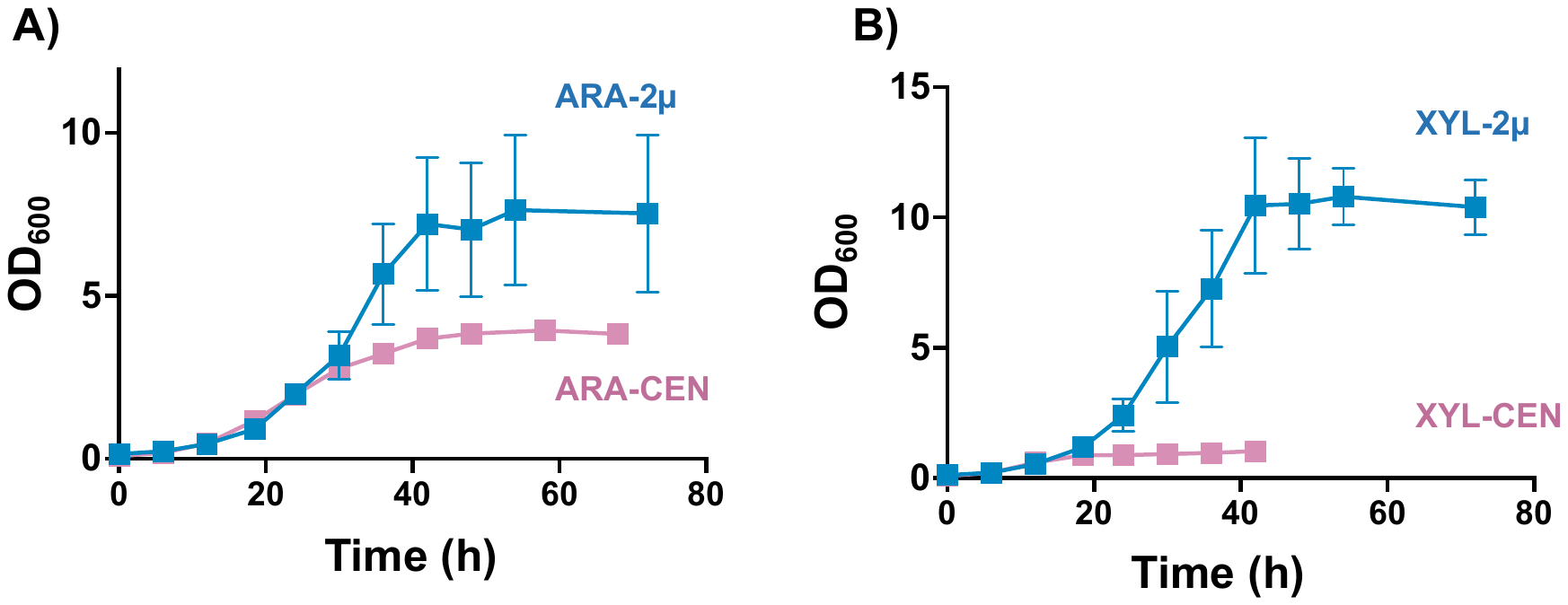

**Figure S3.** Effect of copy number on growth of REG-strain on A) arabinose and B) xylose. For both carbon sources, the REG strain carrying the respective metabolic plasmids were pre-grown in SC media with sucrose (2%) and subcultured in 2YP with either arabinose (2%) or xylose (2%). The metabolic cassettes for arabinose (*araB-araA-araD*) and xylose (*xylA-xks1*) were cloned in pRS413 for maintaining low-copy number and pRS423 for maintaining high copy number. The growth-rate was calculated in the exponential phase using GraphPad Prism software.

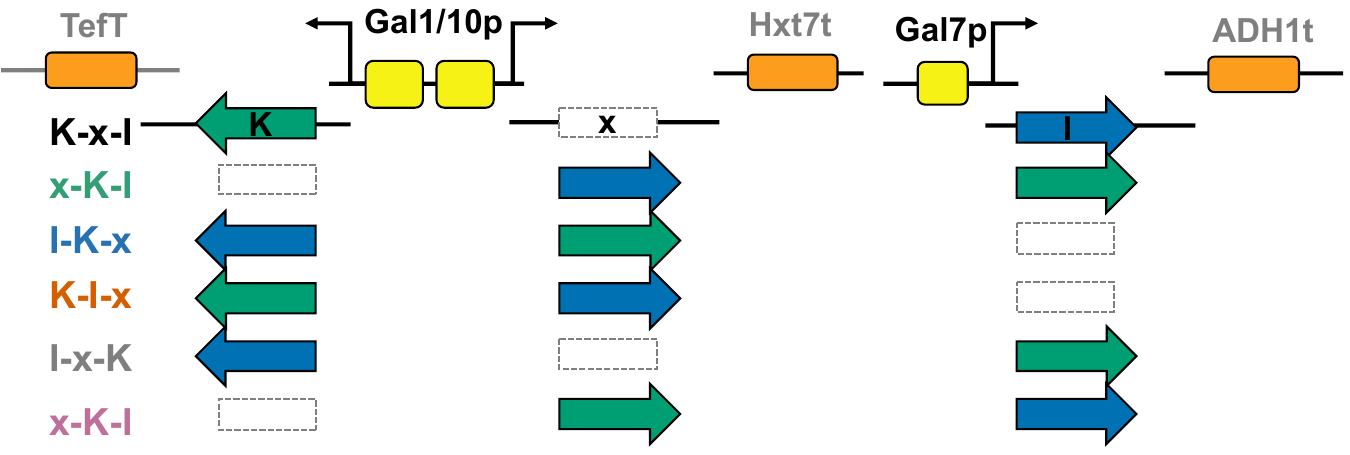

**A)**

**B)**

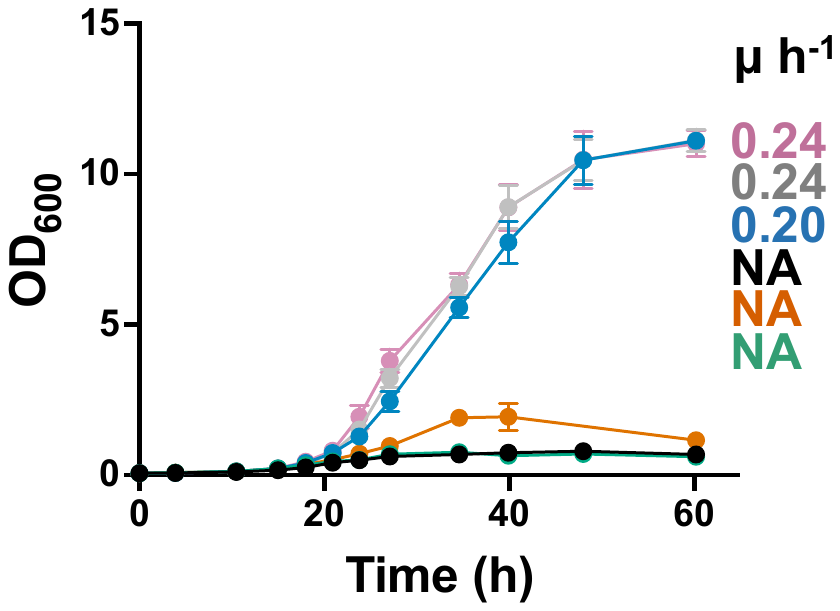

**Figure S4.** A) Schematic of six combinations for xylose metabolic expression constructs generated via promoter-gene swap. The DNA fragments (promoter, gene, and terminator) were assembled in different combinations using NEB-HiFi assembly in the pRS423 background. Yeast xylokinase (K) is represented in green arrows, *Piromyces sp.* XYLA*3 xylose isomerase (I) is represented in blue arrows. Rectangle with broken lines represents lack of any gene (x). B) Comparison of growth performance of different promoter-gene combinations. The culture grown on SC media with sucrose (2%) was inoculated in 2YP with xylose (2%). The growth rate was calculated in the exponential phase using GraphPad Prism software. NA = not determined.

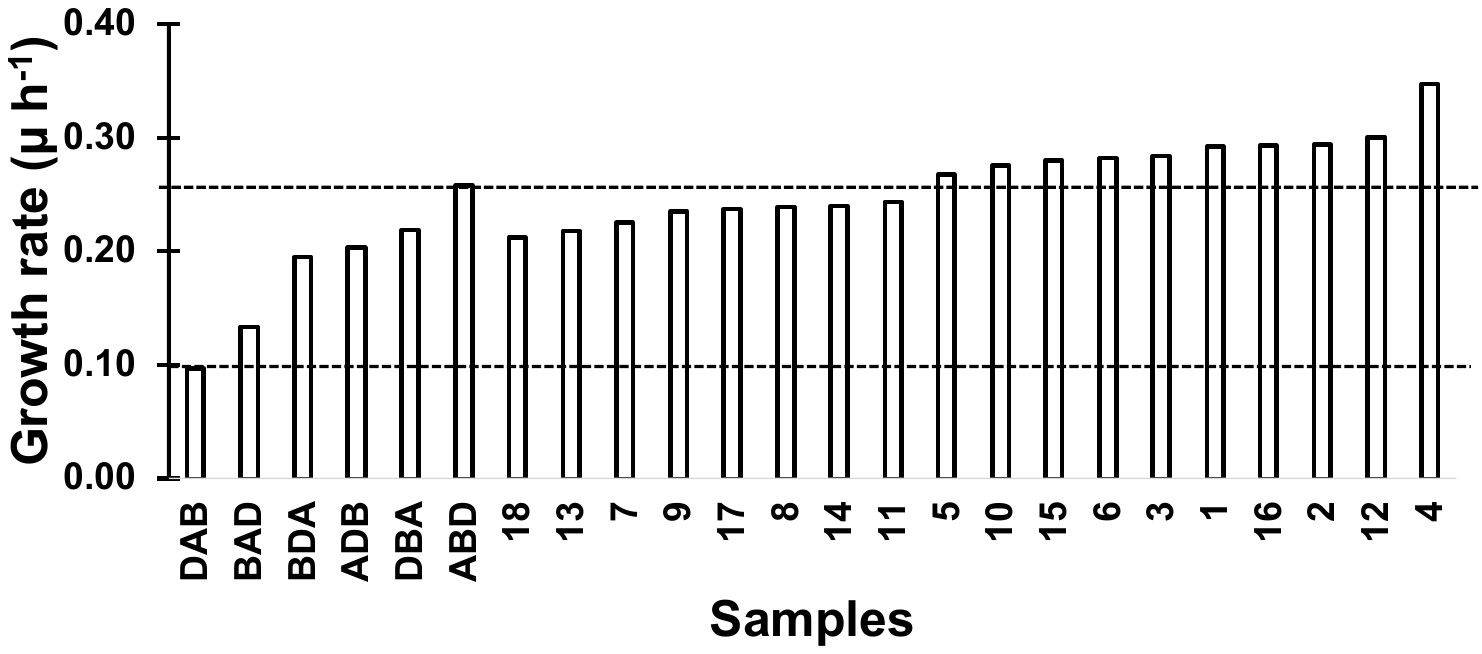

**Figure S5.** Growth rate of colonies screened after library enrichment in SC+Ara. At the end of enrichment cycle (passage # 15), the library pool was plated on SC agar with arabinose (2%). To test the growth-rate of 24 randomly picked colonies, culture was pre-grown on SC with sucrose (2%) and inoculated in SC with arabinose (2%). The growth rate was calculated in the exponential phase using GraphPad Prism software. The screen was performed in a single experiment to identify potential variants for detailed study. Among the top 10 performers we randomly selected 4 variants (3, 4, 12, and 16) for plasmid sequencing. The arabinose cassettes from the variants were amplified and re-cloned into pRS413 and re-transformed in REG-strain for growth-rate determination. This was done to avoid the influence of mutations accumulated in the plasmid backbone / genome due to passaging.

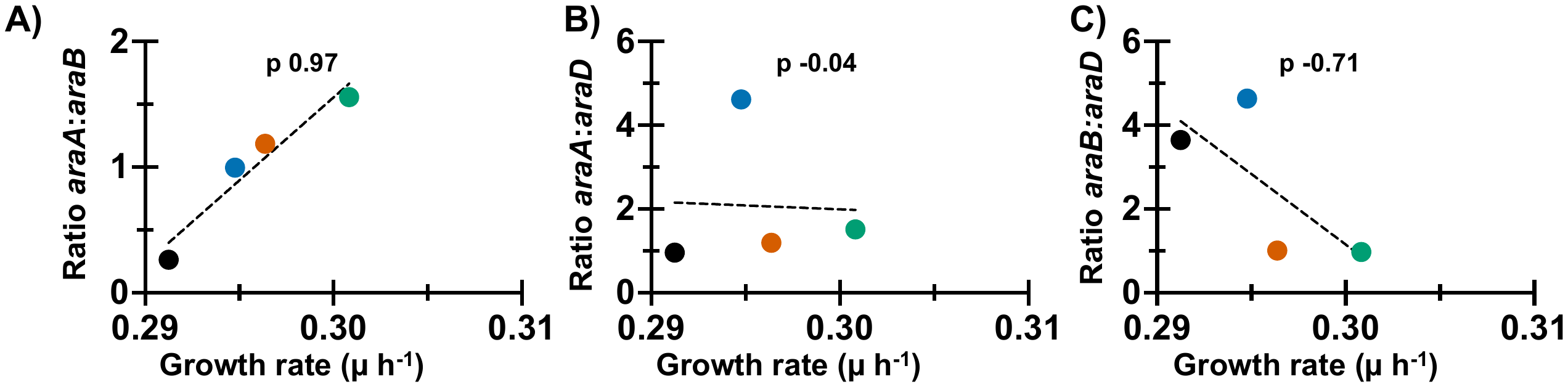

**Figure S6**. Correlation between growth rate and gene expression levels of **A)** *araA*:*araB*, **B)** *araA:araD,* and **C)** *araB:araD* on plasmids isolated from best performing strains after directed evolution. Values indicated are Pearson’s correlation coefficient.
